# Nucleic Acid Adductomics – the Next Generation of Adductomics for Assessing Environmental Health Risk

**DOI:** 10.1101/2022.03.24.485617

**Authors:** Marcus S. Cooke, Yuan-Jhe Chang, Yet-Ran Chen, Chiung-Wen Hu, Mu-Rong Chao

**Author notes:** Address correspondence to: M. S. Cooke,; C. W. Hu,; M. R. Chao,. Marcus S. Cooke, and Yuan-Jhe Chang contributed equally to this work.

## Abstract

The exposome describes the totality of internal and external environmental exposures, across the life course. Components of the exposome have been linked to an increased risk of various, major diseases. To identify the precise nature, and size, of risk, in this complex mixture of exposures, powerful tools are needed to link exposure, cellular consequences, and health/disease. The most biologically informative biomarkers of the exposome should, to varying extents, reflect the dose of the exposure on the body or target organ(s), a subsequent effect on the biological system and, ideally, possess a role in disease. Modification of nucleic acids (NA) is a key consequence of environmental exposures, and while cellular DNA adductomics aims to evaluate the totality to DNA modifications in the genome, an approach which encompasses modifications of all nucleic acids, would be far more comprehensive, and therefore informative. To address this, we propose a cellular and urinary NA adductomics approach for the assessment of both DNA and RNA modifications, including modified (2’-deoxy)ribonucleosides (2’dN/rN), modified nucleobases (nB), plus: DNA-DNA, RNA-RNA, DNA-RNA, DNA-protein, and RNA-protein crosslinks (DDCL, RRCL, DRCL, DPCL, and RPCL, respectively). Supporting the feasibility of this approach, we presented preliminary, proof-of-principle results, which revealed the presence of over 1,000 modified NA moieties, and at least six types of NA modifications, in a representative, pooled urine from healthy subjects, including modified 2’-dN, modified rN, modified nB, DRCL, RRCL and RPCL, many of which were novel/unexpected. We suggest that NA adductomics will provide a more comprehensive approach to the study of nucleic acid modifications, which will facilitate a range of advances, including the identification of novel, unexpected modifications e.g., RNA-RNA, and DNA-RNA crosslinks; key modifications associated with mutagenesis; agent-specific mechanisms; and adductome signatures of key environmental agents, leading to the dissection of the exposome, and its role in human health/disease, across the life course.

## Introduction

The Environment, in its broadest sense, represents 80-90% of the risk of developing cancer, and degenerative diseases^1, 2^. Specifically, this risk of disease is driven by components of the exposome i.e., all the internal and external, environmental factors to which humans are exposed, across the life course, including diet, air pollution, social interactions, and lifestyle choices ^3^ (Figure 1). Exposome-related health issues therefore represent a significant public health issue ^4^. In the US, one in every two adults (162 million people) reported having a chronic illness (cardiovascular disease, cancer, cardiopulmonary disease, asthma, diabetes, arthritis) ^5^. These conditions have a worldwide impact on the public health of nations, and represent a huge economic burden e.g., in 2018, chronic illnesses cost the US ~$1.1 trillion, ~6% of the nation’s GDP ^6^.

**Figure 1.**
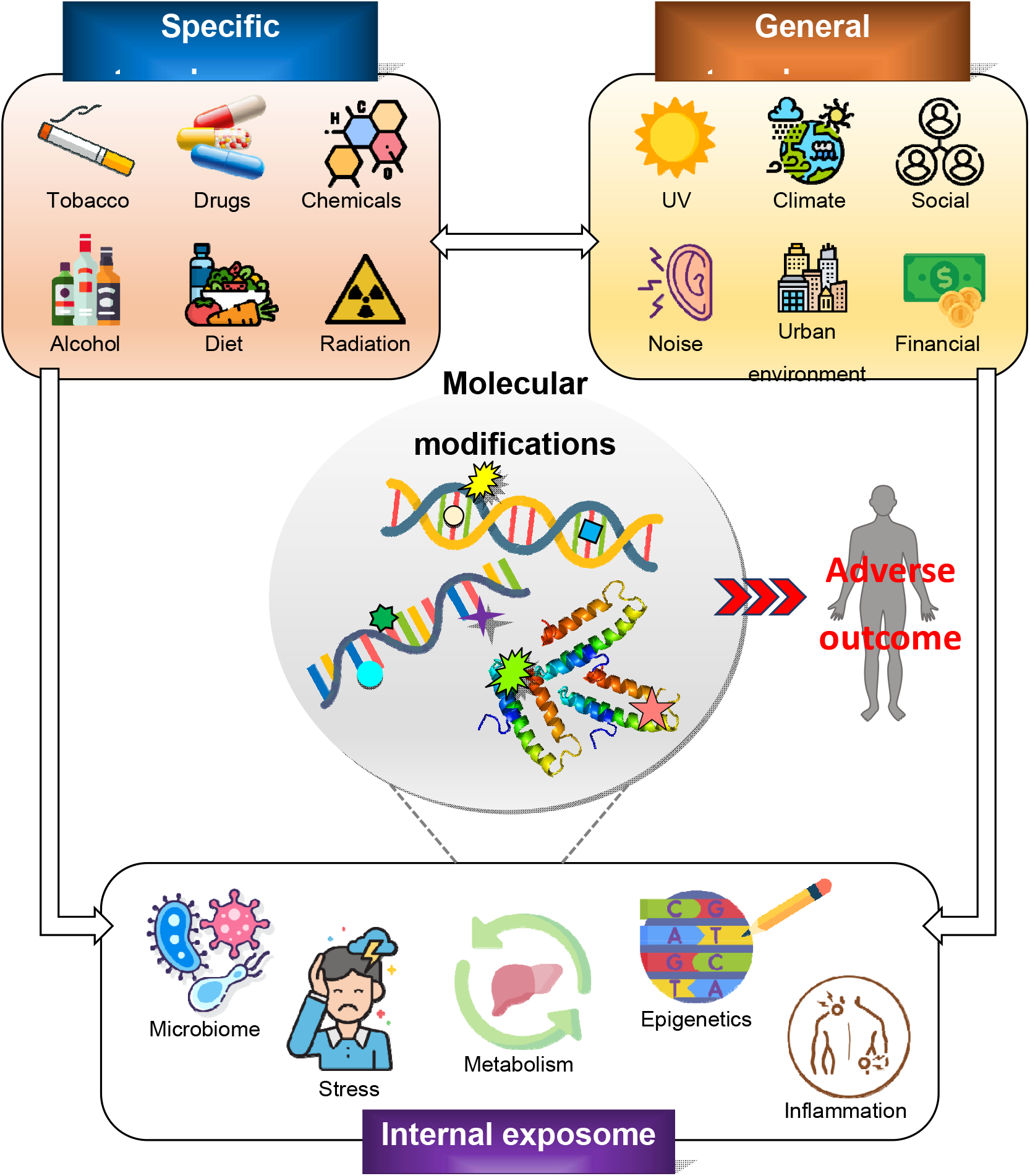
Role of the exposome in the generation of cellular molecular modifications. The exposome (encompassing both external and internal exposomes) represents a wide, and diverse range of exposures, that may occur across the life course. Species arising from the exposome have the potential to damage, or otherwise modify, DNA, RNA, proteins. These modified molecules may result in adverse outcomes for the cell, and lead to disease in the organism. Example exposures from the exposome are illustrated.

Interactions between the exposome and the genome impact individual health, and contribute to the risk of disease ^7^. However, the exposome represents a complex mix of exposures, processes, and factors, and therefore, to identify the precise nature, and size, of risk, powerful tools are needed to link exposure, cellular consequences, and health/disease ^8^. Considerable effort is focused currently on methodology for estimating the specific and general external exposomes (the concepts of which are illustrated Figure 1) i.e., the origins of exposure lie outside the body. From a health perspective, the focus is on the agents which reach the body’s cells, and may then lead to disease ^9^. In this regard, it is the internal exposome which is of primary interest as it represents the total target (which could be a particular cell type, organ, or whole-body burden) dose of physico-chemical agents in the body, irrespective of their source or how they are derived (Figure 1) ^8, 9^ (we have added ‘physico-’ to the definition to include agents such as radiation). A biomarker is a measurable substance in an organism whose presence is indicative of some phenomenon such as disease, infection, or environmental exposure. Therefore, biomarkers of the internal exposome e.g., modifications of cellular biomolecules (Figure 1) may be particularly informative biologically, as they may represent the internal dose, effect(s) on the biological system, and possess a role in disease ^10^. On this basis, a major goal is to use biomarkers of the internal exposome to (i) identify and (ii) act as proxies for, key health-modifying elements of the exposome without the need for detailed measurements of the external environment ^8^. In this Perspective, we have summarized some of the recent advances in DNA adductomics, an approach which aims to describe the totality of DNA adducts derived from environmental and dietary genotoxicants and endogenously produced electrophiles ^11^, and hence inform on the internal exposome. We then indicated how DNA adductomics may advance to become more comprehensive in the breadth of detected adducts, and hence better inform on both exposures and their biological consequences.

Arising from endogenous and exogenous origins, there is continual exposure to agents, and processes, which damage ^12^, or otherwise modify nucleic acids [DNA, RNA, and the (d)NTP pools; NA] ^13^, and their associated proteins (Figure 1). As discussed elsewhere, whether a particular alteration of NA is referred to as damage or a modification depends upon the modification itself, and context ^14^. Modifications induced intentionally by normal cellular process, such as epigenetic programming (e.g., DNA methylation), may be viewed as modifications, whereas those formed unintentionally are damage ^14^, although a grey area may exist in which context is key to interpretation ^15^.

To date most focus has been upon damage to DNA as this impacts cellular function ^16^, and play a critical role in the pathogenesis of, arguably, all major human diseases e.g., cancer, neurodegeneration, and cardiovascular disease, plus aging ^17^. However, there is increasing evidence that damage to, or modification of, RNA (and the ribonucleotide pool) also affects cell function, and play a role in pathogenesis ^18–20^.

### Origins and the multiplicity of nucleic acid modifications

Processes which alter NA can have exogenous and/or endogenous origins. An example of an endogenous origin is the “leakage” of electrons from normal cellular metabolism, which may lead to the generation of reactive oxygen species, and can damage NA, resulting in background levels of oxidatively modified NA ^21^. Another example is DNA methylation, where specialised enzymes add or remove methyl groups from DNA nucleobases to control gene expression, chromosomal structure, and stability ^22^. In contrast, environmental stressors have exogenous origins, and while the damage that they cause to NA can be specific, and hence characteristic, to a particular stressor e.g., cyclobutane pyrimidine dimers (from UV radiation ^23^), 8,9-dihydro-8-(N7-guanyl)-9-hydroxyaflatoxin B1 (from dietary aflatoxin B1 ^24^), 7-(2’-deoxyadenosin-N^6^-yl)aristolactam I (from Aristolochic acid ^25^), they may also impact endogenous processes, and influence levels of endogenously-derived NA modifications. For example, exposure to UV or ionising radiation, and metabolism of certain xenobiotics may all lead to the generation of free electrons, impact redox homeodynamics, and increase levels of oxidatively modified NA above baseline levels e.g., 8-oxo-7,8-dihydro-2’-deoxyguanosine and 8-oxo-7,8-dihydroguanosine, from DNA and RNA, respectively ^26^.Similarly, some exogenous stressors e.g., benzo(a)pyrene ^27^, polychlorinated biphenyls, methylmercury, and organochlorine pesticides ^28^, can influence methylation and other epigenetic processes Consequently, there is some overlap in the types of damage arising from both endogenous and exogenous sources, and while such forms of damage are therefore not, individually, unique signatures of a particular stressor, they can perhaps be used, in conjunction with other forms of damage, to create stressor-specific signatures, particularly when multiple forms of damage are taken into account, or identify the precise mechanisms by which a stressor may act ^27^. Of the processes which induce DNA damage, oxidative stress is a prime example of the potential to induce a multiplicity of different forms of damage. Recently defined as a hallmark of environmental insult ^29^, oxidative stress generates over 24 DNA nucleobase (nB) products, and the total number of adducts exceeds 100, when 2-deoxyribose (2-dR), and phosphate backbone modifications are considered ^17^, and this does not include DNA-DNA and DNA-protein crosslinks, or the adducts derived from secondary processes, such as lipid peroxidation ^30^. These numbers are increased further, when RNA-derived modifications, and epigenetic modifications of DNA and RNA are also included, bringing the total number of potential modifications/adducts into the high hundreds, if not greater.

Most of the literature describes the targeted analysis of DNA damage, rather than RNA, and the measurement of a single, or a few adducts ^31^. These cannot assess the spectrum of NA adducts present in the cell, which gives rise to several weaknesses: e.g., potential biomarkers of exposure may be excluded, it is impossible to identify modifications which act in concert to increase disease risk, and it precludes the identification of yet-to-be-discovered modifications with a key role in disease. Indeed, there is growing evidence that RNA modifications are involved in a growing number of human diseases^19, 32, 33^ (see also the MODOMICS website for a comprehensive review, https://iimcb.genesilico.pl/modomics/), and RNA modifications have a broad range of functions, including RNA processing, splicing, polyadenylation, editing, structure, stability, localization, translation initiation, and gene expression regulation ^34^. To date, approximately 151 modified RNA nucleobase residues have been identified, increasing to ~334 as nucleoside and nucleotide forms are included ^35–40^ however little is known about their impacts upon cellular function ^13^. Other types of NA-associated adducts also have significant biological consequences e.g., DNA-DNA, and DNA-protein crosslinks (DDCL, DPCL) interfere with DNA replication and transcription ^41^, RNA-protein crosslinks (RPCL) can modulate, or stabilize, RNA, interfering with function ^42^ [DNA-RNA crosslinks (DRCL), and RNA-RNA crosslinks (RRCL) may also occur]. Therefore, non-targeted, adductomic approaches (i.e., studying the totality of adducts ^43^) are required to comprehensively detect the range of potential NA adducts, simultaneously.

### Cellular and urinary DNA adductomics - the path towards NA adductomics

Kanaly *et al.* ^44^ first proposed a cellular DNA adductomic strategy to simultaneously measure multiple DNA adducts using liquid chromatography-triple quadrupole tandem mass spectrometry (LC-QqQ-MS/MS) with electrospray ionization (ESI), operating in the selected reaction monitoring (SRM) mode. Due to the labile nature of the 2-dN glycosidic bond, this method detects a number of contiguous SRM of [M+H]^+^ to [M+H-116]^+^ transitions, in which 116 Da corresponds to the neutral loss of the 2-dR moiety. Lately, high resolution MS (HRMS) has also been adopted for DNA adductomics analysis, for example, the hybrid quadrupole-linear ion trap orbitrap MS (Q-LIT-OT-MS). The mass of the ions detected by OT can be measured with ppm accuracy (≤ 1-5 ppm mass error). This high mass accuracy significantly improves/enhances confidence in DNA adduct identification, compared to the nominal mass accuracy obtained by QqQ-MS/MS. The advent of DNA adductomics by HRMS has led to the proposal of a “top-down” approach, by which patterns of adducts may be used to trace, and identify the originating exposure source (summarised in Figure 2).

**Figure 2.**
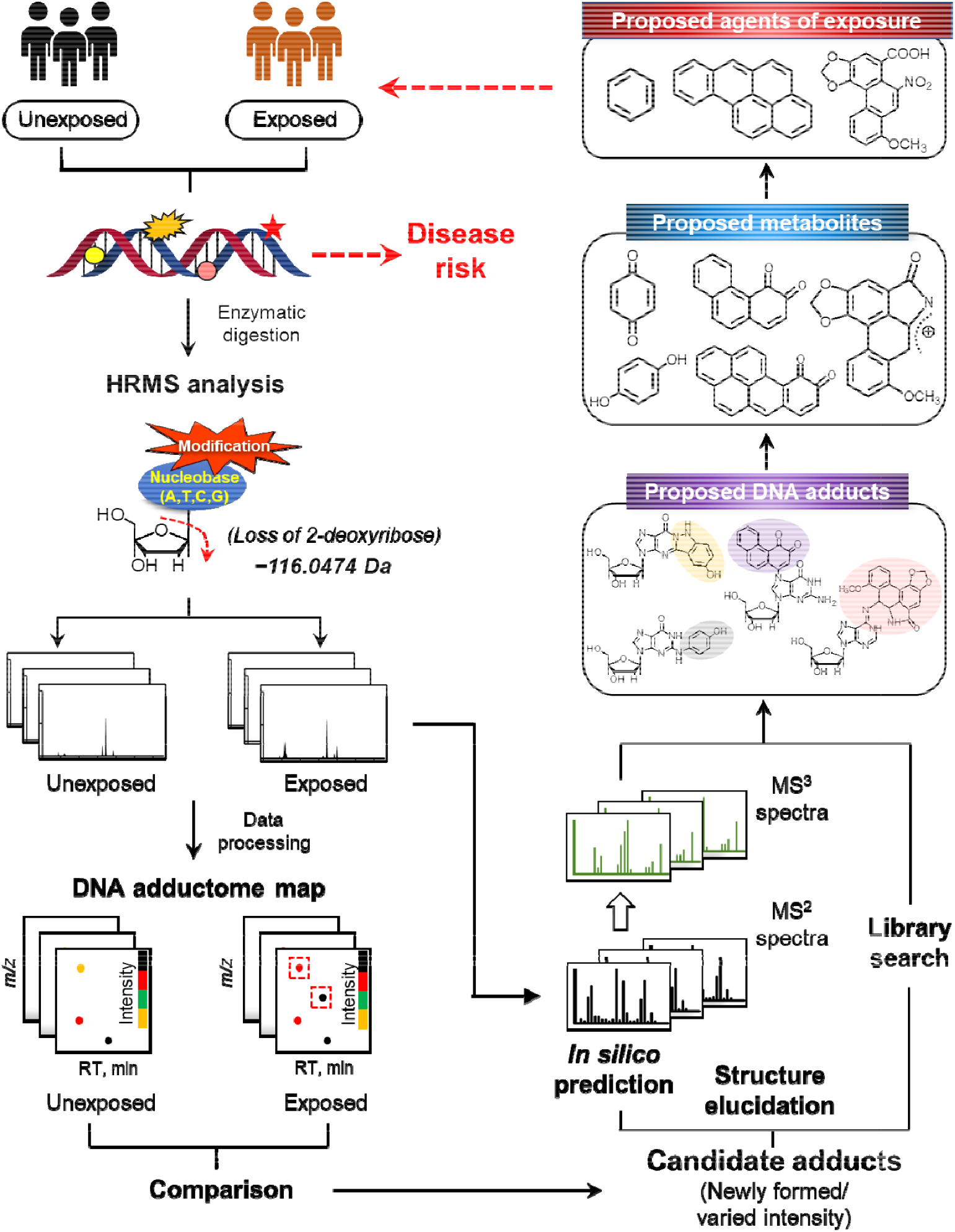
The concept of a top-down approach for HRMS-based DNA adductomics for linking human, environmental carcinogen exposure, and cancer risk. Cellular DNA adductome maps are developed, using HRMS, from exposed and unexposed individuals. DNA adductome maps are compared, and potential candidate adducts, reflective of exposure, are short-listed. Expected (known) adducts are identified from characteristics held within an DNA adductome library ^108^, and unexpected (unknown) adducts are identified by *in silico* prediction. Precursor reactive species/metabolites are identified from the profile of adducts formed, and the responsible environmental agents, to which the individuals were exposure, are then proposed.

Previously, we reported a cellular DNA adductomics approach ^45^, then a HRMS approach ^46^, and broadened the range of targets to include DDCL (i.e., crosslinkomics) ^47^. Recently, RNA adductomics was reported, which uses neutral loss targeting of the [M+H]^+^ > [M+H-132]^+^ transitions, in which 132 Da corresponds to the neutral loss of the 2-ribose (R) ^48^. This revealed the potential for genotoxins to cause analogous forms of damage to both DNA, and RNA ^48^ emphasising further the importance of NA, more broadly, as a target for genotoxins.

### Cellular nucleic acid adductomics

To date, there are no adductomics methods for the combined detection of DNA, and RNA modifications, let alone combined with DDCL, RRCL, DRCL, DPCL, and RPCL. We therefore propose a strategy to perform NA adductomics, as the most comprehensive approach for the study of nucleic acid modifications, to date. In support of this strategy, we illustrated that different NA modifications produce their own specific fragmentation patterns in collision-induced dissociation (CID; Figure 3) by a HRMS (i.e., Q-LIT-OT-MS). The experimental details and method applied are given in the Supplementary information. For example: modified 2’-dN tend to lose dR, resulting in the product ion of [M+H-dR]^+^; modified ribonucleosides (rN) tend to lose ribose (R), resulting in the product ion of [M+H-R]^+^ or [M+H-MeR]^+^, respectively. Modified nB produce specific product ions including [B+H]^+^ and [B+H-NH_3_]^+^; DDCL generate product ions of [M+H-2dR]^+^, and [M+H-dR]^+^; RRCL have product ions of [M+H-2R]^+^ and [M+H-R]^+^; DRCL have product ions of [M+H-dR]^+^, [M+H-R]^+^ and [M+H-dR-R]^+^; DPCL give [M+H-dR]^+^ [M+H-Cys]^+^ and [M+H-dR-Cys]^+^; RPCL generate [M+H-R]^+^ [M+H-Cys]^+^ and [M+H-R-Cys]^+^. The above exact mass of neutral loss (Da) or product ions (m/z) monitored by HRMS are provided in the Supplementary Table S1. It is worth noting that adductomics is not limited solely to adducts formed via damaging reactions i.e., the products of which are unintentional. For example, intentional products, such as those arising from endogenous processes, such as epigenetic modifications ^15^, are also detected by NA adductomics. As noted above, while these are not sufficiently characteristic to identify a particular stressor, changes in their levels are likely to inform, at least in part, on the mode of action of the stressor. The goal is to evaluate the maximal number of modifications, in a single analysis run, which is critical to most effectively characterizing the role of NA adducts in disease risk identification, and prevention.

**Figure. 3.**
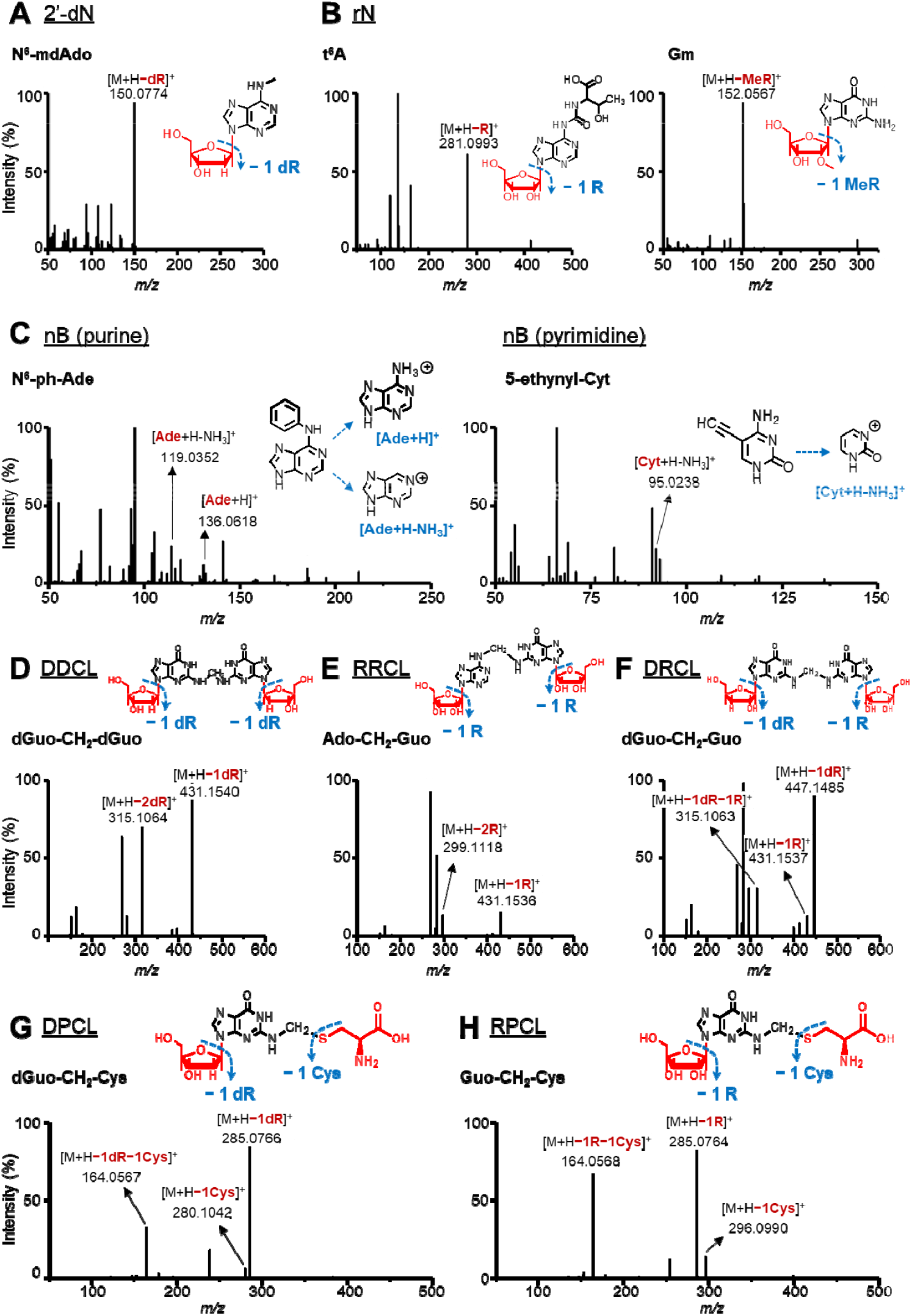
Fragmentation patterns of (A) modified 2’-dN, (B) modified rN, (C) modified nB, (D) DDCL, (E) RRCL, (F) DRCL, (G) DPCL, and (H) RPCL, by LC-Q-LIT-OT-MS with CID fragmentation mode. Test solutions were prepared by dissolving various commercially available standards in deionized water, including modified 2’-dN, modified rN, modified nB. For the DDCL, RRCL, DPCL and RPCL, these were synthesized by incubating native 2’-deoxynucleosides (i.e., dGuo, dAdo, dCyd and dThd), ribonucleosides (i.e., Guo, Ado, Cyd and Urd), and/or cysteine, with formaldehyde ^47^. The fragmentation patterns were obtained using a LC-Q-LIT-OT-MS equipped with a heated electrospray ionization source, operated in positive ESI-CID mode. All the informative, and specific product ions for each type of NA modification were generated in MS^2^. The experimental details are provided in the Supplementary information.

While cellular DNA, and hence NA, adductomics have multiple, potential applications ^49^, these are likely to be most informative when applied human population studies. However, there have been few reports of cellular DNA adductomics analysis of human tissue ^44, 50^, let alone populations. Human tissue represents a severe analysis challenge due to the need for high sensitivity to detect multiple adducts ^51^. Recent advances in linear ion trap, and Orbitrap instruments offer improved sensitivity ^31^, but the availability of such instrumentation is limited, not least due to cost. Furthermore, cellular DNA adductomics requires a significant amount of DNA, typically 50-500 μg, depending upon adduct frequency ^52, 53^, which is rather more than that needed for targeted analyses ^31^. Obtaining tissue is necessarily invasive, can be challenging logistically, adds additional ethics review board scrutiny, may discourage volunteer participation ^54, 55^, and limits access to vulnerable individuals (e.g., the young, or elderly). Isolating DNA from exfoliated epithelial cells (e.g., from the urogenital tract or buccal cavity) represents a potential way forward, and a means to circumvent some of these issues ^56^, albeit while restricting the investigation to a particular organ (which may or may not be a bad thing). In contrast, biomonitoring NA damage products in urine offers numerous advantages over cellular NA^30^. Indeed, we recently reported urinary DNA adductomics, which circumvents many of the issues associated with cellular DNA adductomics ^57^, and we propose that the same strengths apply to urinary NA adductomics.

### Biomonitoring NA adducts in urine

The presence of DNA adducts in urine is generally considered to be the consequence of DNA repair ^58, 59^ i.e., base excision repair (BER) is the source of nB adducts ^60^, and global genome-, and transcriptional coupled-nucleotide excision repair (GG-NER, and TCR-NER, respectively), results in 2’-dN adducts ^61–64^ (Figure 4). More specific evidence for this comes from the reported urinary presence of 2’-dN adducts, such as (5’R)-, (5’S)-8,5-cyclo-2’deoxyadenosines ^65–67^, and 3- (2-deoxy-β-d-erythro-pentofuranosyl)pyrimio[1,2-α]purin-10(3H)-one ^68, 69^, all of which appear to be repaired by NER ^69–71^. Additionally, sanitization of the (2’-deoxy)ribonucleotide pools represents another logical source of urinary 2’-dN (and rN) adducts (Figure 4), a proposal we have maintained for some time ^72^. Spontaneous depurination may also be a contributor to urinary adduct levels ^73^ e.g., unstable N7-guanine adducts, which have been reported to occur dose-dependently in the urine of rodents, and which are in proportion to levels in the liver (reviewed in ^74^).The urinary adductome will be reflective of the body burden of adducts, given an on-going exposure to environmental agents which may induced DNA damage, and the usual steady-state repair^75^. While it may be argued that these repaired adducts are unlikely to be the ones specifically responsible for inducing mutations, or other adverse cellular events, they do reflect the risk of mutations occurring i.e., the larger the adductome burden, the greater the risk of mutation occurring ^76^. Furthermore, defects in the post-adduct excision steps of DNA repair may also have detrimental consequences for the cell^77^

**Figure 4.**
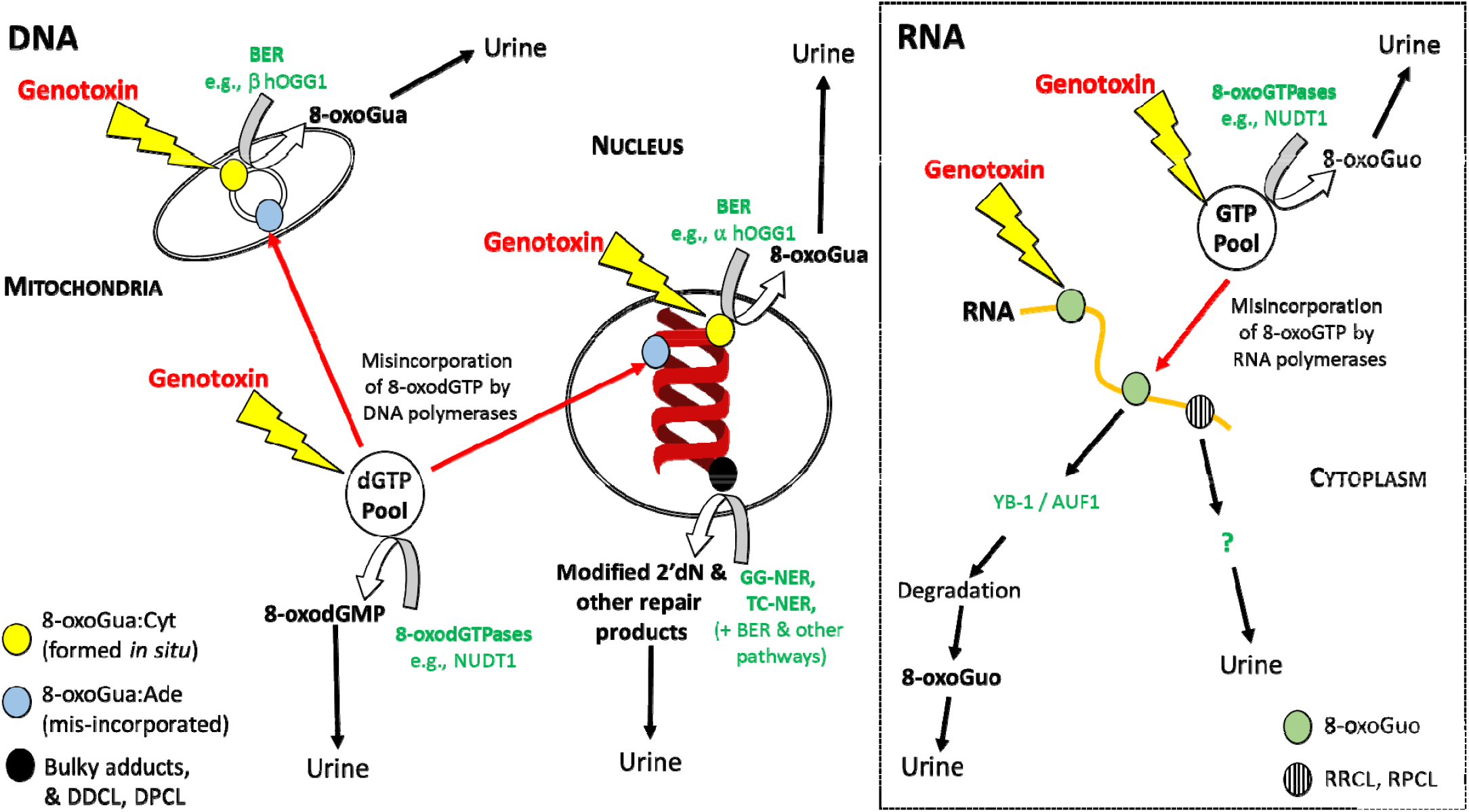
Representative sources of NA modifications in urine. Broadly, base excision repair (BER) of DNA leads to the presence of modified nucleobases in urine, and there is some evidence that urinary, modified 2’-deoxyribonucleosides are derived from the activity of global genome, and/or transcription-coupled, nucleotide excision repair (NER). The activity of enzymes, such as the Nudix hydrolases (e.g., NUDT1), which hydrolyse modified (2’-d)NTPs to NMPs, may also contribute to urinary levels of modified (2’-deoxy)ribonucleosides. The figure uses oxidatively modified DNA and RNA to illustrate examples of the processes. The repair products of DNA-DNA crosslinks (DDCL), DNA-protein crosslinks (DPCL), RNA-RNA crosslinks (RRCL), and RNA-protein crosslinks (RPCL) are less well defined.

There is a growing number of reports where targeted approaches have been used to successfully detect specific, environmentally-induced DNA adducts in urine. Examples of urinary adducts include: 3-(4-dihydroxyphenyl)adenine, following benzene exposure of mice ^78^; aflatoxin N7-guanine, and 8,9-dihydro-8-(2,6-diamino-4-oxo-3,4-dihydropyrimid-5-yl-formamido)-9-hydroxy-aflatoxin B1, following exposure of rats to aflatoxin ^79^; N7-(1-hydroxy-3-buten-2-yl)guanine following exposure of mice to 1,3-butadiene (BD) ^80^, and occupationally- or recreationally- (tobacco smoke) exposed humans ^81^; N7-(1-hydroxy-3-buten-2-yl) guanine adducts were also detected in the same human urine samples from the previous citation ^80^, and *bis*-N7-guanine DNA-DNA crosslinks in mice exposed to BD ^82^. This work, using targeted approaches, is beginning to successfully link exposure to environmental agents to specific urinary DNA adducts. Taken together, these findings are highly supportive of the potential of urinary NA adducts to lead to signature NA adductome profiles of key environmental exposures.

The literature describes a growing number of reports describing processes for the removal of RNA-derived adducts, and these are supported by the presence of RNA products in urine (reviewed in ^32^; Figure 4). The processes for the repair of crosslinks (NA-NA, and NA-protein) are more complex, and less well understood than for mono-adducts. DDCL are repaired by processes that involves NER^83^, with the potential involvement of BER^84, 85^. It has been suggested that the crosslink is removed in the context of an oligonucleotide, which is subsequently digested down to a smaller, lesion-containing product, prior to excretion into the urine ^64^. There are reports of multiple mechanisms for cells to cope with DNA-protein crosslinks ^86^, which may be broadly defined as: direct hydrolysis of the covalent linkage between protein and DNA; excision of the lesion-containing DNA fragment; and proteolysis of the protein component ^87^. Given these possibilities, the post-repair processing of the DNA-protein adduct itself is less clear than for DDCL (and for RRCL, and RPCL, even less so) ^88^. Nevertheless, unless the adduct is reversed, some form of adduct-derived product will remain, which needs to be removed, and excreted (as evidenced from our data, presented here).

Urine has numerous benefits as a matrix in which to study biomarkers. It is obtained non-invasively, easily collected, transported, and stored, with low biological hazard. Furthermore, no pre-processing is needed prior to storage, small volumes are required for analysis (typically less than 500 μL), and adduct stability is maintained for >15 y at −80 °C (in the case of the representative lesion, 8-oxodGuo, at least)^89^, allowing the use of previously banked samples ^90, 91^. Urine sample workup is generally simpler than for DNA. It is a well-established matrix in which to study biomarkers of exposure ^92^, including a wide variety of DNA adducts e.g., oxidized nucleic acid products (including RNA) ^93, 94^, *N*-nitroso-derived DNA adducts ^95^, alkylated purines ^96^, and aristolochic acid-derived DNA adducts ^97^, illustrating the breadth of adducts in urine, which parallels that in NA. However, as noted above, such targeted analyses severely restrict the amount of information that can be gained. Applying adductomics to the study of urine therefore represents an innovative, and exciting means to non-invasively assess the totality of NA modifications, and therefore exposures.

While cellular and urinary DNA adductomics aim to assess the totality of DNA adducts in the genome, these approaches are not entirely comprehensive for genomic damage, as they have excluded RNA modifications (i.e., damage, and intentional modifications, such as epigenetic changes), and other classes of adducts (e.g., crosslinks), all of which have downstream consequences for the cell, and are consequently implicated in pathogenesis ^32, 41, 42, 98–100^. We propose that the NA adductome represents a superior, functional, multi-faceted biomarker of the interaction between exposome, and genome. Furthermore, comprehensively assessing the totality of damage, to all NA moieties, is critical to sensitively, and specifically, quantify, and characterize exposures, and define the mechanistic role, of nucleic acid damage in cancer risk, and prevention. The strengths of this approach include consideration of: (i) the differences in the adducts, or modifications, formed, which arises from the physicochemical nature of the NA moiety, e.g., due to abundance, conformation (double-, single-stranded, or free nucleoside/nucleobase), physical associations (e.g., with proteins, or metal ions), location (nucleus, cytoplasm, proximity to mitochondria, and propensity to modification (e.g., RNA is more easily oxidised than DNA) ^101, 102^ (and probably more easily damaged generally); (ii) the variety of downstream effects that modified nucleic acids can cause, and their relevance to pathogenesis, e.g., mutations (DNA), epigenetic changes (DNA/RNA), error-containing proteins, and altered protein synthesis (RNA) ^32^; and (iii) the ability to perform these assessments non-invasively, via urine.

### Evidence for the feasibility of urinary NA adductomics

Cellular DNA adductomics of biological samples is becoming increasingly well established (recent publications include ^48, 103–105^, and references contained in ^11^). However, as noted above, it possesses three key weaknesses, it is neither: (i) entirely comprehensive for DNA adducts, (ii) nor does not detect RNA modifications (which are of importance in the eukaryotic response to environmental stresses, and pathogenesis ^106^), (iii) nor is it easily amenable to the study of human populations. We have addressed these issues, with the development of cellular and urinary NA adductomics, extending our cellular ^45^, and urinary ^57^ DNA adductomics assays to include modified 2’-dN, nB, rN, DDCL, RRCL, DRCL, DPCL, and RPCL. Urine is the ideal matrix in which to assess the NA adductome, as it contains the totality of adducts, from the repair of cells in the entire body, producing levels significantly higher than in individual tissues, and reflecting the totality of exposures. We therefore propose a NA adductomics approach, using a LC-Q-LIT-OT-MS with CID fragmentation, to achieve the detection of the widest, and most comprehensive, range of nucleic acid modifications. To support the feasibility of our proposed urinary NA adductomics approach, our preliminary, proof-of-principle results revealed that at least six types of NA modifications were present in the urine of healthy subjects (Figure 5), including modified 2’-dN, modified rN, modified nB, DRCL, RRCL and RPCL. DDCL and DPCL were not detected in the urine, from healthy human subjects. Of the NA modifications detected, three out of 61 modified 2’-dN, 16 out of 265 modified rN, six out of 325 modified nB were fully identified, and confirmed using commercially available standards (summarized in Table S2). The detailed LC-Q-LIT-OT-MS analysis is provided in the Supplementary information. Among these fully identified NA modifications, it is worth noting that these include a number of novel NA modifications, not previously identified in the human urine, such as N^4^-methyl-2’-deoxycytidine (N^4^-mdCyd). At present, we cannot identify whether they derive from exogenous, or endogenous sources (or both). As expected, several well-known NA modifications were also successfully detected, such as, 8-oxodGuo and 5-methyl-2’-deoxycytidine (5-mdCyd; see Figure 5A), N^6^-methyladenine (m^6^A) and N^6^, 2⍰-O-dimethyladenosine (m^6^Am; see Figure 5B), together with N7-methylguanine (N7-mGua), N3-methyladenine (N3-mAde) and 8-oxoGua (Figure 5C). In addition to the modified purine nucleobases (Figure 5C), a number of modified pyrimidine nucleobases were also observed (> 400 adducts; data not shown). Regarding the NA-associated crosslinks, excitingly, one DRCL (Figure 5D) and six RRCL (Figure 5E) were identified, which we propose to be AP apurinic/apyrimidinic (AP) site-derived crosslinks ^107^ (DRCL: Ion 1 = 2’-deoxyadenosine-AP (dAdo-AP); RRCL: Ion 2 = 2’-O-methylguanosine-AP (Gm-AP); Ion 3 = 2’-O-methyluridine-AP (Um-AP); Ion 4 = deazaguanosine-AP (dzGuo-AP); Ion 5 = adenosine-AP (Ado-AP); Ion 6 = 2’-O-methyladenosine-AP (Am-AP); Ion 7 = adenosine-di-AP (Ado-Di-AP). Moreover, a novel RPCL (Ion 8) was newly identified, which we propose is adenosine-CH_2_-cysteine (Ado-CH_2_-Cys). These assumptions were supported by the presence of their protonated precursor ions [M + H]^+^, as well as their specific fragmentation features. The proposed chemical structures, and product ion spectra of NA-associated crosslinks are provided in Supplementary Figure S1-S3, and Table S3. In addition to the above NA modifications, we detected numerous other NA modifications in human urine, but these are yet to be fully identified (e.g., Figures 5A-5C). The structure elucidation of these NA modifications requires a searchable mass spectral database, such as the one currently under development for DNA adductomics ^108^. In total, we detected over 1,000 modified nucleic acid moieties in urine, giving the first indication of the significant breadth of modifications present in the cellular, and urinary, adductomes.

**Figure 5.**
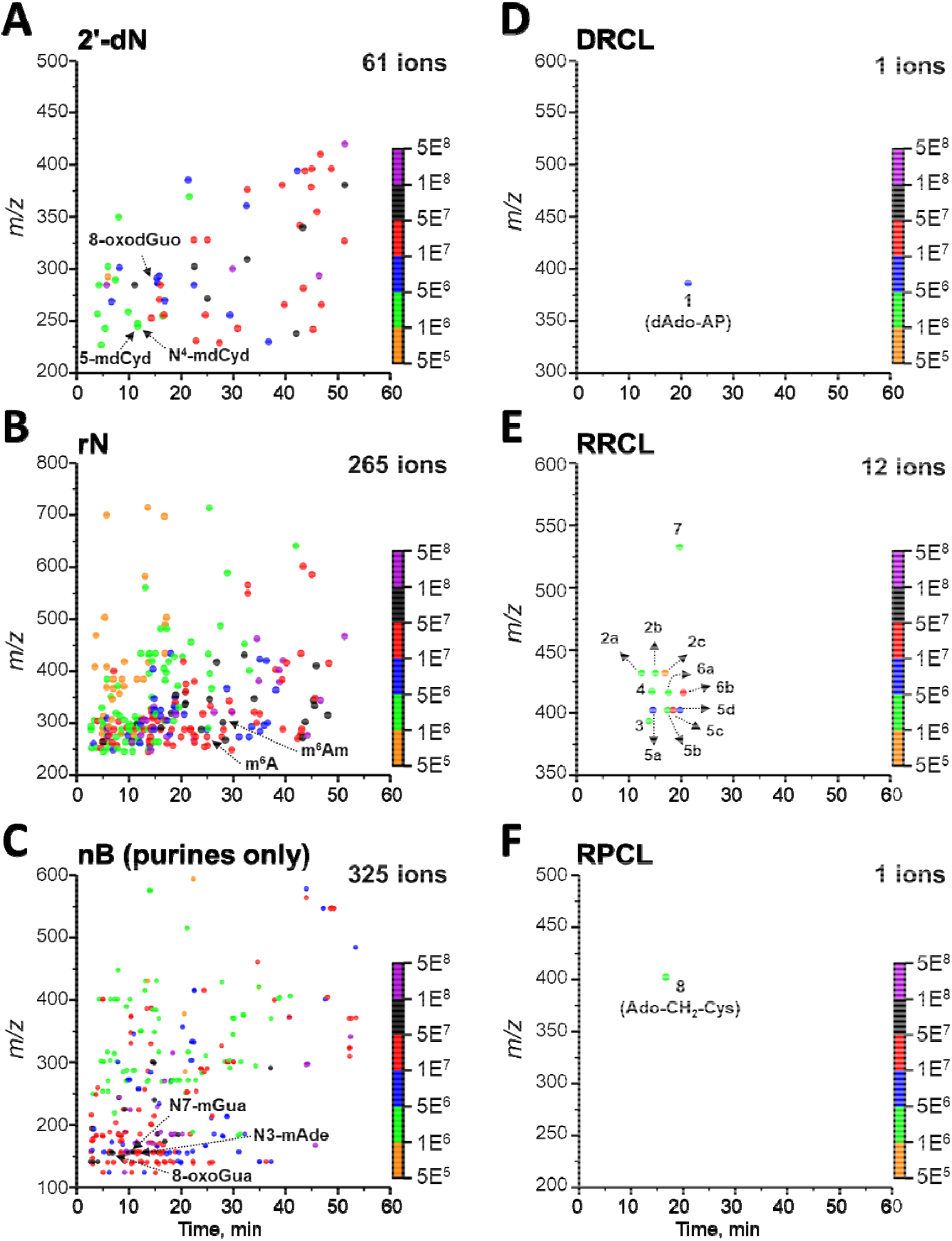
Proof-of-principle: successful generation of NA adductome maps for (A) 2’-dN adducts, (B) rN adducts, (C) nB adducts (modified purines), (D) DRCL, (E) RRCL and (F) RPCL, from a pooled urine derived from six healthy subjects, by LC-Q-LIT-OT-MS with CID fragmentation. Data are reported as accurate mass-to-charge ratios (m/z; Y axis, left) with retention times (RT; X axis) and associated peak intensities (spot color; Y axis, right), using OriginPro. Eight NA-associated crosslinks (Ions 1-8) were newly identified, and the corresponding isomers of RRCL were further labelled with letter “a” to “d”. Additionally, over 300 ions were noted corresponding to pyrimidine nB (data not shown). The identification of each type of NA modification was achieved by the observation of their protonated precursor ions [M + H]^+^, and their specific fragmentation features in MS^2^. The detailed HRMS monitoring parameters are given in the Supplementary information.

HRMS-omics analysis always generates a massive dataset of product ion spectra. In the present study, the above NA modifications were characterized, and identified one by one using an ion-map function of Thermo Scientific Xcalibur software (ver. 4.1, Thermo Fisher Scientific Inc., USA) combined with extensive manual inspection, which is a complicated and extremely time-consuming process. This laborious process highlights the need of dedicated, post-data analysis software, which can characterize, and identify all the specific fragmentation features derived from all types of NA adducts. To date, post-data analysis software is only available for the DNA adductomics^51, 109^.

Despite of the lack of post-data processing software for NA adductomics, we demonstrate the ability to detect novel, and/or unexpected NA modifications, such as the modified 2’-dN, rN, DRCL, RRCL and RPCL noted here, and the several crosslinks we reported previously ^47^. Combined, this represents a major strength of NA adductomics, and offers a means to fully characterise exposures based upon the spectra of adducts that they induce, together with providing a better understanding of the modifications which lead to mutagenicity.

### Challenges

There are a number of challenges to performing NA adductomics successfully, using HRMS, these include: (1) the need of high-throughput post-data processing software, as the data sets generated are massive and complex, consisting of various features derived from different types of NA modifications. (2) The establishment of an NA adductome database would assist in adduct identification. Unlike proteomics or metabolomics, a NA adductome database is not available yet, although a DNA adductome database is in preparation ^108^. This would allow searches based on exact mass, fragmentation pattern, and chemical structures, as well as offering visual representations of the 3D structure, for the identification of NA modifications. (3) Harmonization of LC-HRMS-based NA adductomics approaches. NA adductomics can be performed using a range of HRMS instruments. Many pre-analytical, analytical, and post-analytical factors can impact NA adduct detection, identification, and quantification. In the near future, strategies are needed to account for these differences including data normalization, removing matrix effects and ion suppression, or using pooled quality control samples and internal standards, for harmonizing cross-platform and cross-laboratory data collected from untargeted HRMS-based NA adductome studies. For example, a reference standardization protocol has been recently proposed for harmonizing large-scale metabolomics data collected across different studies or different HRMS methods ^110^. Analogous issues have been addressed for the measurement of cellular and urinary nucleic biomarkers of oxidative stress via the formation of international consortia, such as the European Standards Committee on Oxidative DNA Damage, the European Standards Committee on Urinary Lesion Analysis, and the European Comet Assay Validation Group ^111^. We propose that an equivalent consortium, encompassing all that NA adductomics represents, would be the beginning of a strategy towards addressing the above challenges.

### Future perspectives, and conclusions

We suggest that the NA adductomic approach to biomonitoring environmental exposure will offer a valuable means to contribute to evaluating the exposome, and its impact upon health/disease, across the life-course. For the first time, through the detection of over 1,000 modified NA moieties, we illustrate the scale of the NA adductome, further emphasising the need, and potential benefits, of the NA adductomic approach. The impact of this methodology extends beyond cancer, to any disease for which there is an environmental component, and for which our approach will facilitate Adductome-Wide Association Studies (AWAS; an untargeted, agnostic, hypothesis-generating approach for exploring nucleic acid/protein modifications associated with health outcomes, complementary to GWAS, Epigenome-WAS, and Exposome-WAS). As show in Figure 6, we propose applying our approach to the creation of a library of NA adductome maps, consisting of the integrative and informative NA adductome profiles for relevant, environmental agents. More broadly, this can be combined with the study of associations between DNA, and RNA modifications, and the relationships between environment-induced chemical modifications, and epigenomic modification (which we term the “NA adductome network”). Our approach also has further utility for mechanistic studies, e.g., for more comprehensively identifying the processes responsible for genotoxicity; elucidating the spectrum of physiologically relevant NA modifications that are targeted by discrete repair pathways; improving our understanding of how environmental agents impact epigenetics; and performing more detailed exploration of the relationship between NA adductome signatures, mutational signatures ^52, 112, 113^, epigenome/epitranscriptome signatures ^34, 114^, and metabolome signatures ^115^. Combined, this will provide the most comprehensive, multi-adductomic characterizations of environmental agents to date, vital to identifying exposomic factors associated with disease, their mechanisms, and with potential utility in disease detection, prevention, and risk assessment.

**Figure 6.**
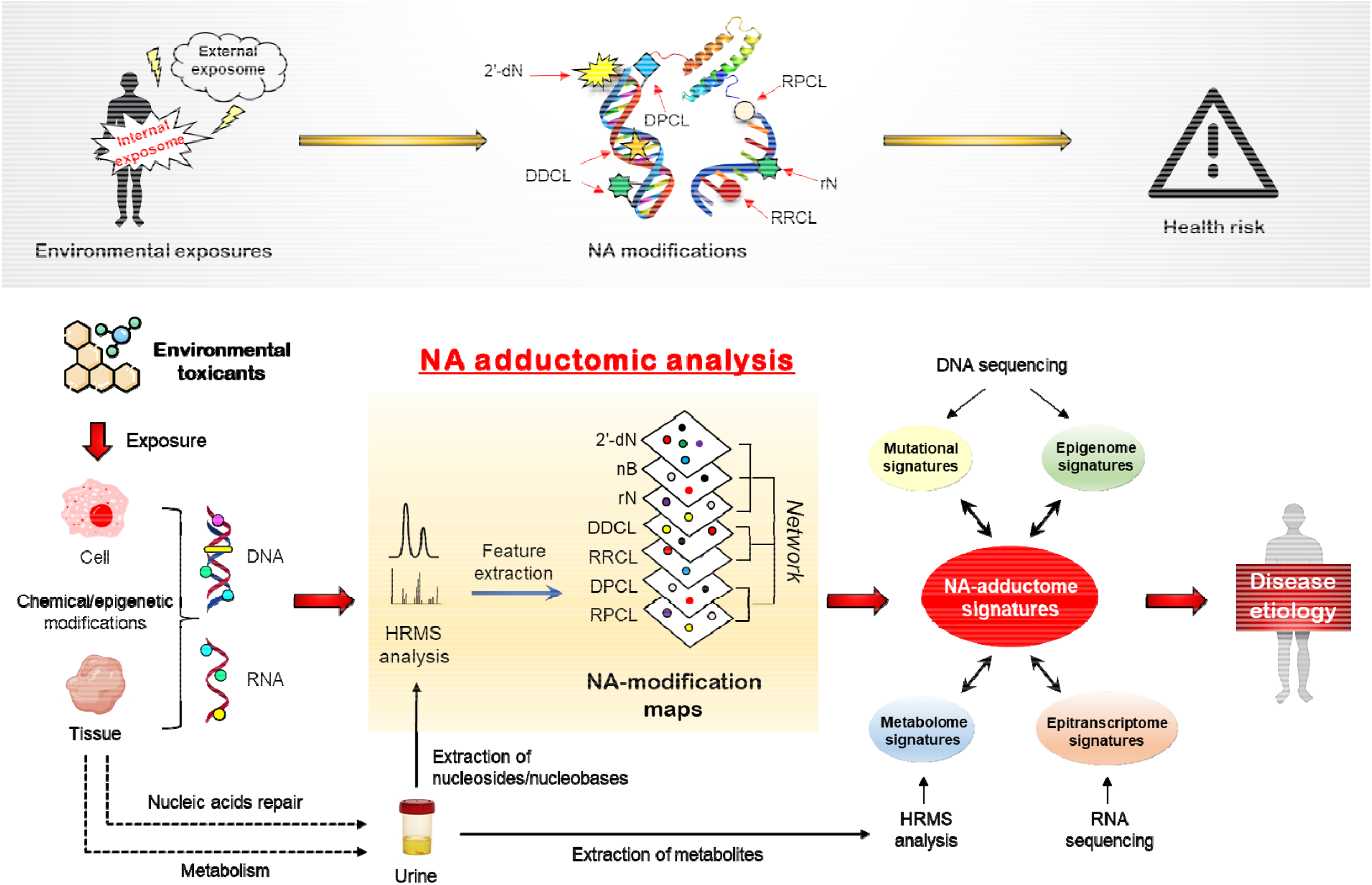
Illustration of how Adductome-Wide Association Studies (AWAS) may be used to explore the environmental causes of cancer. The proposed approach may be used to investigate the link between exposure to environmental agents, and health risk. **Top panel.** The premise is that the exposome results in modification of all types of nucleic acids [DNA, RNA, (d)NTPs], which is associated with an increased risk of developing major diseases, such as cancer. **Main figure.** *In vitro* and *in vivo* models are exposed to a panel of environmental toxicants, individually, and cellular and urinary NA adductome maps are created. These are used to develop a library of NA adductome maps and a network of inter-related NA modifications. Associations are then investigated between NA adductomic “signatures” of specific exposures and mutational, epigenomic, metabolomic, and epitranscriptomic which, together, may elucidate the underlying etiology and/or mechanism(s) of disease.

## Acknowledgements

The research reported in this publication was supported, in part, by the National Institute of Environmental Health Sciences of the National Institutes of Health under award number R01ES030557 (to MSC and C-WH), and R15ES027196 (MSC). The content is solely the responsibility of the authors and does not necessarily represent the official views of the National Institutes of Health. This work was also supported by the Ministry of Science and Technology, Taiwan (grant numbers MOST 109-2314-B-040-018-MY3 to M-RC; MOST 110-2628-B-040-002 to C-WH).

Declaration of competing financial interests: The authors declare they have no actual or potential competing financial interests.

## CRediT author statement

Marcus S. Cooke: Conceptualisation, Writing - Original draft preparation, Reviewing and Editing; Yuan-Jhe Chang: Investigation, Data curation; Yet-Ran Chen: Methodology, Data curation; Chiung-Wen Hu: Conceptualisation, Methodology, Writing - Reviewing and Editing; and Mu-Rong Chao: Conceptualisation, Methodology, Data curation, Writing - Reviewing and Editing.

## Notes

### Competing Interest Statement

The authors have declared no competing interest.

