## Supplemental information for "Nucleic Acid Adductomics – the Next Generation of Adductomics for Assessing Environmental Health Risk"

**Experimental Information**

1. **Preparation of various NA crosslinks induced by formaldehyde**

Formaldehyde solution was prepared by diluting an aqueous solution of formaldehyde (37% w/v) with deionized water to yield a concentration of 250 mM as working solution. To produce various formaldehyde-induced NA crosslinks, 80 μL of formaldehyde solution (250 mM) and 280 μL of 0.1 M phosphate buffer (pH 7.0) were incubated with 20 μL of 50 mM each of native 2'-deoxynucleosides (i.e., dGuo, dAdo, dCyd and dThd), ribonucleosides (i.e., Guo, Ado, Cyd and Urd), and/or cysteine, at 37 °C for 72 h ^1^.

**2. Urine sample collection and preparation**

This study was approved by the Institutional Review Board of Chung Shan Medical University Hospital in Taiwan (CSMUH No: CS2-21079). All the participants were adults. Written, informed consent was obtained from the participants. Urine samples were obtained from six adults aged from 22-45 y, and pooled to create a single sample. Prior to analysis, the pooled urine was filtered through a centrifugal filter (Nanosep®, 3K, Omega™ membrane, Pall Laboratory, USA), and was then diluted twice with deionized water, containing four stable isotope-labelled internal standards (^15^N_5_-8-oxodGuo, d_4_-N^6^-(2-hydroxyethyl)-2'-deoxyadenosine, d_3_-N2-(deoxyguanosin-8-yl)-2-amino-3,8-dimethylimidazo[4,5-f]quinoxaline and d_3_-5-hydroxymethyl-2'-deoxycytidine), ready for analysis.

**3. NA adductomics analysis by LC-HRMS**

*3.1. LC separation and hybrid quadrupole-linear ion trap-orbitrap MS (Q-LIT-OT-MS) analysis*

A previously validated LC-HRMS method was used ^2^. The formaldehyde-induced DNA adducts were separated by a Thermo Fisher Scientific Vanquish UHPLC system (Waltham, MA, USA) with an Inertsil ODS-3 C18 column (150 × 2.1 mm, i.d., 5 μm) from GL Sciences (Tokyo, Japan). The mobile phase consisted of 1 mM aqueous ammonium acetate solution as mobile phase A, and 95% methanol (v/v) containing 0.1% (v/v) formic acid as mobile phase B.

An Orbitrap Fusion Lumos Tribrid Mass Spectrometer (Thermo Fisher Scientific) equipped with a heated electrospray ionization (HESI) source was used and operated in positive ESI mode. The source voltage was 3.5 kV. The gas setting of sheath gas, aux gas and sweep gas were 35, 7 and 0 arbitrary units, respectively. The vaporizer and ion transfer tube temperatures were optimized to be 200 ℃. Non-targeted NA modifications analysis was achieved using the data-dependent acquisition mode (i.e., dd-MS^2^). Initially, the MS^1^ full scan was performed iteratively in orbitrap with a resolution of 60,000 over a mass range of *m/z* 100 - 1000. The instrument was run in top speed mode with a cycle time of 2.5 seconds. The intensity greater than 25,000 from the MS^1^ were automatically selected and fragmented in the linear ion trap (MS^2^ fragmentation by collision-induced dissociation, CID) at a collision energy of 30%.

The MS^2^ spectra were acquired in the orbitrap with a resolution of 15,000. A dynamic exclusion of 6 seconds was used. A targeted exclusion list, derived from the background ions of solvent blank injection, was applied in the MS^2^ fragmentation to improve the sensitivity of the detection of NA modifications.

*3.2. Generation of the featured peak lists by ion mapping*.

Raw data files generated from the dd-MS^2^ scan were processed directly using the Xcalibur software version 4.1 (Thermo Scientific) ^3^. Initially, the *m/z* values of each specific fragmentation feature were inputted sequentially to generate the corresponding ion maps (i.e., neutral loss map or parent map) in the Qual Browser interface. The neutral loss maps were used to screen for the 2'-dN adducts, rN adducts and NA crosslinks, while the parent maps were used to screen for the nB adducts (modified purines or pyrimidines). The neutral loss map was generated based on the detection and display of all precursor ions that had a neutral loss of specific mass (as listed in Table S1; e.g., neutral loss of 116.0474 Da for the moiety of one deoxyribose). For the parent map, it was generated based on detection and display of all precursor ions that had specific fragment ions (product ions) corresponding to four different nucleobases, such as, *m/z* 151.0494 ([Gua]^•+^), 135.0545 ([Ade]^•+^), 111.0433 ([Cyt]^•+^) and 126.0429 ([Thy]^•+^), as shown in Table S1. The mass tolerance was ± 0.01 *m/z* for both neutral loss map and parent map. Both maps consisted of parent *m/z* (X-axis), product *m/z* (Y-axis), and intensity (Z-axis) with an informative mass spectrum, displaying all precursor ions detected. The detected precursor ions were then subsequently exported to Microsoft Excel for developing the peak lists. The output peak list consisted of *m/z* and intensity of the precursor ions. The retention time of precursor ions in the peak list were assigned manually by inspection of raw data. The false-positive signals were also excluded by manual inspection of the fragments observed in the MS^2^ spectra. Finally, the detected ion signals were converted to a visual, colored NA adductome map, which consisted of RT (X-axis), *m/z* (Y-axis), and peak intensity (spot color), using OriginPro 2016 Sr2 software (MA, USA).

**Table S1.** Parameters used for generating both neutral loss map or parent map.

| NA modifications | Exact mass of neutral loss (Da) or  product ion (*m/z*) applied | Description of parameters |
| --- | --- | --- |
| Modified 2'-dN | -116.0474 | Neutral loss of dR |
| Modified rN | -132.0423 | Neutral loss of R |
|  | -146.0579 | Neutral loss of MeR |
| DDCL | -232.0948 | Neutral loss of 2 dR |
| RRCL | -264.0845 | Neutral loss of 2 R |
|  | -292.1158 | Neutral loss of 2 MeR |
|  | -278.1002 | Neutral loss of R + MeR |
| DRCL | -248.0896 | Neutral loss of dR + R |
|  | -262.1053 | Neutral loss of dR + MeR |
| DPCL | -237.0671 | Neutral loss of dR + Cys |
| RPCL | -253.0620 | Neutral loss of R + Cys |
|  | -267.0777 | Neutral loss of MeR + Cys |
| Modified nB | 151.0494 | Product ion *m/z* of [Gua]^•+^ |
|  | 152.0567 | Product ion *m/z* of [Gua + H]^+^ |
|  | 135.0301 | Product ion *m/z* of [Gua + H − NH_3_]^+^ |
|  | 135.0545 | Product ion *m/z* of [Ade]^•+^ |
|  | 136.0618 | Product ion *m/z* of [Ade + H]^+^ |
|  | 119.0352 | Product ion *m/z* of [Ade + H − NH_3_]^+^ |
|  | 111.0433 | Product ion *m/z* of [Cyt]^•+^ |
|  | 112.0505 | Product ion *m/z* of [Cyt + H]^+^ |
|  | 95.0240 | Product ion *m/z* of [Cyt + H − NH_3_]^+^ |
|  | 126.0429 | Product ion *m/z* of [Thy]^•+^ |
|  | 127.0502 | Product ion *m/z* of [Thy + H]^+^ |
|  | 110.0237 | Product ion *m/z* of [Thy + H − NH_3_]^+^ |

**Table S2.** Fully identified NA modifications (and Guo) present in the urine of healthy subjects. Each NA modification was confirmed against commercially available reference standards.

| Adduct type | RT (min) | Measured *m/z* ([M+H]^+^) | Full name |
| --- | --- | --- | --- |
| 2'-dN | 11.82 | 242.1134 | 5-methyl-2'-deoxycytidine |
|  | 11.84 | 242.1134 | N^4^-methyl-2'-deoxycytidine |
|  | 15.61 | 284.0991 | 8-oxo-7,8-dihydro-2'-deoxyguanosine |
| rN | 5.87 | 282.1195 | 1-methyladenosine |
|  | 6.77 | 302.0984 | 5-carbamoylmethyluridine |
|  | 8.11 | 298.1146 | 7-methylguanosine |
|  | 8.59 | 274.1033 | 5-hydroxymethylcytidine |
|  | 9.48 | 258.1085 | 5-methylcytidine |
|  | 11.44 | 284.0993 | Guanosine |
|  | 12.70 | 300.0935 | 8-oxo-7,8-dihydroguanosine |
|  | 15.11 | 317.0981 | 5-methoxycarbonylmethyluridine |
|  | 15.35 | 298.1146 | 2'-O-methylguanosine |
|  | 16.35 | 286.1032 | N^4^-acetylcytidine |
|  | 18.04 | 273.1081 | 5-methyl-2'-O-dimethyluridine |
|  | 19.18 | 312.1300 | 2-(dimethylamino)guanosine |
|  | 21.11 | 333.0751 | 5-methoxycarbonylmethyl-2-thiouridine |
|  | 24.11 | 282.1195 | N^6^-methyladenosine |
|  | 27.08 | 413.1413 | N^6^-(N-threonylcarbonyl)adenosine |
|  | 28.32 | 296.1357 | N^6^, 2'-O-dimethyladenosine |
| nB | 8.63 | 126.0663 | 5-methylcytosine |
|  | 8.66 | 126.0662 | N^4^-methylcytosine |
|  | 6.89 | 168.0517 | 8-oxo-7,8-dihydroguanine |
|  | 9.46 | 150.0772 | N3-methyladenine |
|  | 11.57 | 166.0724 | N7-methylguanine |
|  | 18.29 | 180.0879 | N^2^-ethylguanine |

**Table S3.** Proposed identities of the NA-associated crosslinks detected in the urine of healthy subjects, as measured by LC-Q-LIT-OT-MS.

| Adduct type | Ion | Measured *m/z*  ([M+H]^+^) | RT (min) | Theoretical *m/z*  ([M+H]^+^) | Mass error^a^ (ppm) | Confirmation ion  (*m/z*) | Proposed identity |
| --- | --- | --- | --- | --- | --- | --- | --- |
| DRCL | 1 | 384.1520 | 21.56 | 384.1514 | 1.6 | 268.1041 [M+H-1dR]^+^ (or [Ado+H]^+^); 252.1088 [M+H-1R]^+^ (or [dAdo+H]^+^); 136.0616 [M+H-1dR-1R]^+^ (or [Ade+H]^+^) | 2'-deoxyadenosine-AP  (dAdo-AP) |
| RRCL | 2a | 430.1567 | 12.59 | 430.1569 | -0.5 | 298.1146 [M+H-1R]^+^ (or [Gm+H]^+^);  152.0565 [M+H-1R-1MeR]^+^ (or [Gua+H]^+^);  135.0297 [Gua+H-NH_3_]^+^ | 2'-O-methylguanosine-AP  (Gm-AP) |
|  | 2b | 430.1565 | 15.23 |  | -0.9 | 298.1144 [M+H-1R]^+^ (or [Gm+H]^+^);  152.0567 [M+H-1R-1MeR]^+^ (or [Gua+H]^+^);  135.0298 [Gua+H-NH_3_]^+^ |  |
|  | 2c | 430.1566 | 17.13 |  | -0.7 | 298.1146 [M+H-1R]^+^ (or [Gm+H]^+^);  152.0563 [M+H-1R-1MeR]^+^ (or [Gua+H]^+^);  135.0295 [Gua+H-NH_3_]^+^ |  |
|  | 3 | 391.1349 | 14.01 | 391.1347 | 0.5 | 245.0768 [M+H-1MeR]^+^ (or [Urd+H]^+^);  113.0345 [M+H-1R-1MeR]^+^ (or [Ura+H]^+^) | 2'-O-methyluridine-AP  (Um-AP) |
|  | 4 | 415.1459 | 14.59 | 415.1460 | -0.2 | 283.1039 [M+H-1R]^+^ (or [dzGuo+H]^+^);  151.0614 [M+H-1R-2R]^+^ (or [dzGua+H]^+^) | Deazaguanosine-AP  (dzGuo-AP) |
|  | 5a | 400.1459 | 14.80 | 400.1463 | -1.0 | 268.1039 [M+H-1R]^+^ (or [Ado+H]^+^);  136.0617 [M+H-2R]^+^ (or [Ade+H]^+^) | Adenosine-AP  (Ado-AP) |
|  | 5b | 400.1458 | 17.60 |  | -1.2 | 268.1037 [M+H-1R]^+^ (or [Ado+H]^+^);  136.0618 [M+H-2R]^+^ (or [Ade+H]^+^) |  |
|  | 5c | 400.1456 | 18.70 |  | -1.7 | 268.1036 [M+H-1R]^+^ (or [Ado+H]^+^);  136.0619 [M+H-2R]^+^ (or [Ade+H]^+^) |  |
|  | 5d | 400.1459 | 20.01 |  | -1.0 | 268.1039 [M+H-1R]^+^ (or [Ado+H]^+^);  136.0614 [M+H-2R]^+^ (or [Ade+H]^+^) |  |
|  | 6a | 414.1621 | 17.76 | 414.1619 | 0.5 | 282.1200 [M+H-1R]^+^ (or [Am+H]^+^);  268.1040 [M+H-1MeR]^+^ (or [Ado+H]^+^);  136.0617 [M+H-1R-1MeR]^+^ (or [Ade+H]^+^) | 2'-O-methyladenosine-AP  (Am-AP) |
|  | 6b | 414.1620 | 20.64 |  | 0.2 | 282.1201 [M+H-1R]^+^ (or [Am+H]^+^);  268.1039 [M+H-1MeR]^+^ (or [Ado+H]^+^);  136.0619 [M+H-1R-1MeR]^+^ (or [Ade+H]^+^) |  |
|  | 7 | 532.1872 | 19.96 | 532.1885 | -2.4 | 400.1454 [M+H-1R]^+^ (or [Ado-AP+H]^+^); 268.1034 [M+H-2R]^+^ (or [Ado+H]^+^) | Adenosine-di-AP  (Ado-Di-AP) |
| RPCL | 8 | 401.1242 | 16.90 | 401.1238 | 1.0 | 280.1042 [M+H-1Cys]^+^ (or [Ado+C+H]^+^); 269.0820 [M+H-1R]^+^ (or [Ade+Cys+C+H]^+^);  148.0619 [M+H-1Cys-1R]^+^ (or [Ade+C+H]^+^);  136.0622 [Ade+H]^+^ | Adenosine-CH_2_-Cysteine  (Ado-CH_2_-Cys) |

^a^ Mass error: (Measured *m/z* – theoretical *m/z*) / theoretical *m/z* × 10^6^

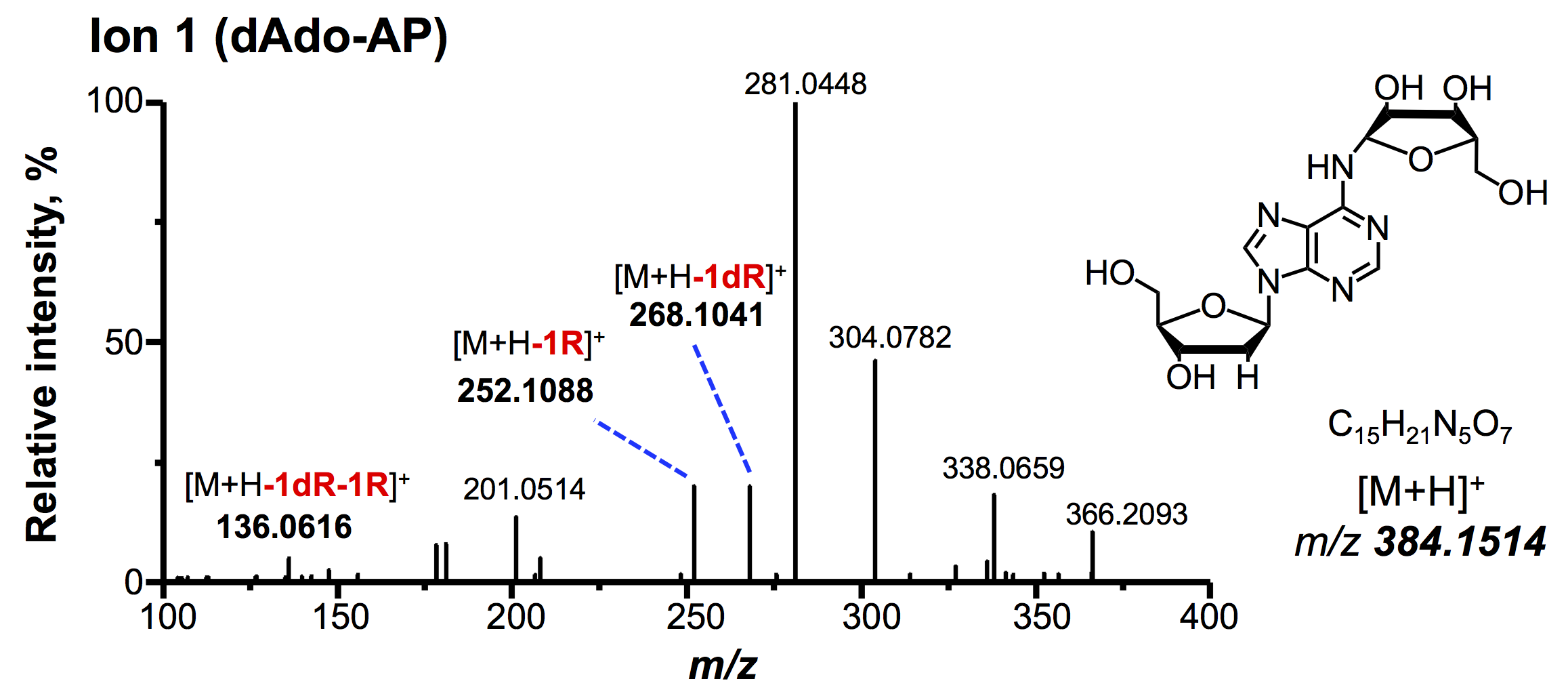

**Figure S1.** Product ion spectra of dAdo-AP (Ion 1), and proposed structure, as determined by LC-Q-LIT-OT-MS with CID fragmentation.

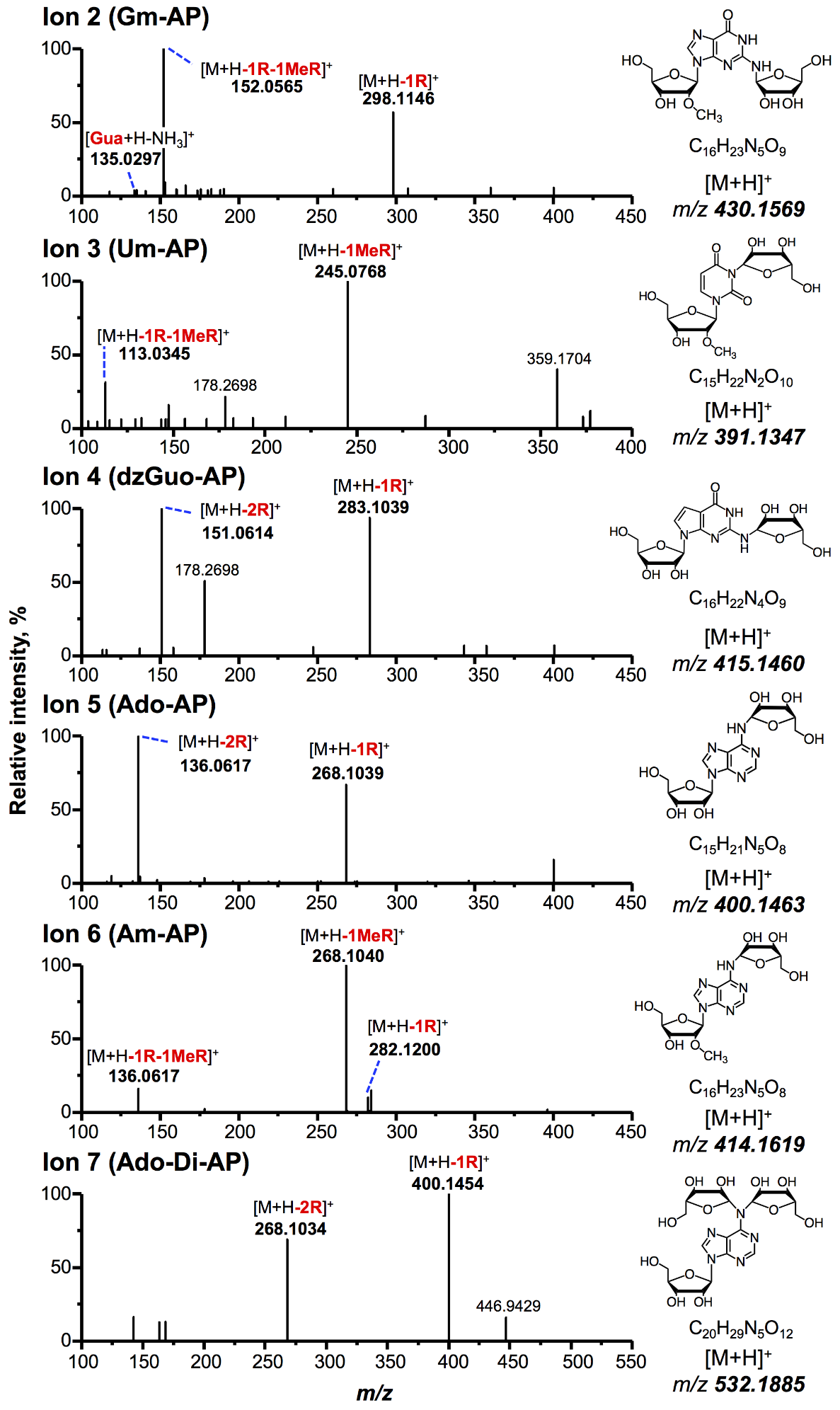

**Figure S2.** Product ion spectra of RRCL crosslinks (Ions 2-7), and proposed structures, as determined by LC-Q-LIT-OT-MS with CID fragmentation.

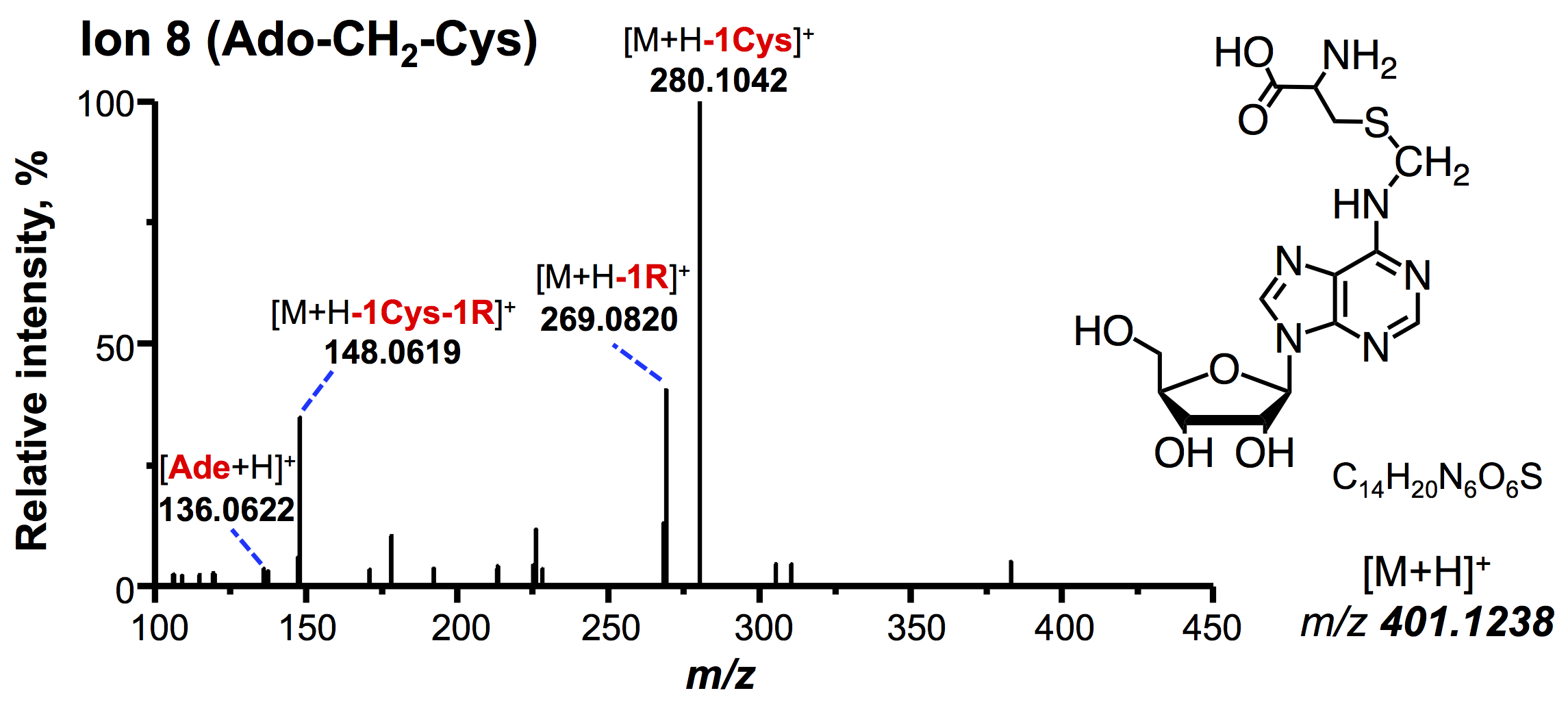

**Figure S3.** Product ion spectra of Ado-CH_2_-Cys (Ion 8), and proposed structure, as determined by LC-Q-LIT-OT-MS with CID fragmentation.
